# Dominant monoallelic variant in the PAK2 gene causes Knobloch syndrome type 2

**DOI:** 10.1101/2020.10.06.328419

**Authors:** Stylianos E. Antonarakis, Ales Holoubek, Melivoia Rapti, Jesse Rademaker, Jenny Meylan, Justyna Iwaszkiewicz, Vincent Zoete, Muhammad Ansar, Christelle Borel, Olivier Menzel, Kateřina Kuželová, Federico A. Santoni

## Abstract

Knobloch syndrome is an autosomal recessive phenotype mainly characterized by retinal detachment and encephalocele caused by biallelic pathogenic variants in the *COL18A1* gene. However, there are patients clinically diagnosed as Knobloch syndrome with unknown molecular etiology not linked to *COL18A1*. We studied an historical pedigree (published in 1998) designated as KNO2 (Knobloch type 2 syndrome with intellectual disability, autistic behavior, retinal degeneration, encephalocele). Whole exome sequencing of the two affected siblings and the normal parents resulted in the identification of a PAK2 non-synonymous substitution p.(Glu435Lys) as a causative variant. The variant was monoallelic and apparently *de novo* in both siblings indicating a likely germline mosaicism in one of the parents; the mosaicism however could not be observed after deep sequencing of blood parental DNA. PAK2 encodes a member of a small group of serine/threonine kinases; these P21-activating kinases (PAKs) are essential in signal transduction and cellular regulation (cytoskeletal dynamics, cell motility, death and survival signaling, and cell cycle progression). Structural analysis of the PAK2 p.(Glu435Lys) variant which is located in the kinase domain of the protein predicts a possible compromise in the kinase activity. Functional analysis of the p.(Glu435Lys) PAK2 variant in transfected HEK293T cells results in a partial loss of the kinase activity. PAK2 has been previously suggested as an autism related gene. Our results show that PAK2 induced phenotypic spectrum is broad and not fully understood. We conclude that the KNO2 syndrome in the studied family is dominant and caused by a deleterious variant in the PAK2 gene.

## Introduction

The sequence analysis and exploration of the human genome has significantly contributed to the identification of the molecular cause of Mendelian disorders. To date, there are 4,587 protein-coding genes linked to Mendelian disorders (OMIM https://www.omim.org/ search of 17-sep-20); there are however approximately 15,000 additional protein-coding genes that are not yet linked to a disorder of a specific trait. On the other hand, the study of Mendelian phenotypes has often revealed different genes responsible for similar phenotypes^1^. An extreme example is the non-specific intellectual disability with mendelian inheritance, where pathogenic variants in more than 700 different genes are responsible for these phenotypes^2^. Knobloch syndrome (KNO1 OMIM 267750) is an autosomal recessive phenotype mainly characterized by retinal detachment and encephalocele caused by biallelic pathogenic variants in the *COL18A1* gene on chromosome 21^3^. The syndrome may present with different extraocular abnormalities: a single umbilical artery, pyloric stenosis, a flat nasal bridge, midface hypoplasia, bilateral epicanthal folds, abnormal lymphatic vessels in the lung, patent ductus arteriosus, cardiac dextroversion, generalized hyperextensibility of the joints, unusual palmar creases, and unilateral duplication of the renal collecting system. Structural brain defects, epilepsy and cognitive abnormalities have also been frequently reported. However, the full phenotypic spectrum is yet to be defined.

There are patients clinically diagnosed as Knobloch syndrome with unknown molecular etiology not linked to *COL18A1* (OMIM 120328). We have studied an historical pedigree from New Zealand first published in 1998^4^, designated as KNO2 (Knobloch type 2 syndrome). Exome sequence of the two affected siblings and the normal parents resulted in the identification of a heterozygous *PAK2* non-synonymous substitution as a causative variant. The variant was monoallelic and apparently de novo in both siblings due to presumed parental mosaicism. PAK2 encodes a member of a small group of serine/threonine kinases; these P21-activating kinases (PAKs) are important in signal transduction and cellular regulation (cytoskeletal dynamics, cell motility, death and survival signaling, and cell cycle progression)^5^. Thus, the KNO2 phenotype due to monoallelic pathogenic variants in *PAK2* has a dominant mode of inheritance.

## Methods

### Family ascertainment and DNA sequencing

DNA samples from the four members of the family (parents and 2 affected siblings) were stored in the DNA bank of the Geneva laboratory since 2003. These DNAs were processed by whole exome sequencing at the Health 2030 Genome Center using Twist Human Core Exome Kit (TWIST Biosciences, San Francisco, CA, USA); sequencing was performed on an Illumina HiSeq4000 platform.

### Computational analysis

The exome-sequencing data were analyzed using a customized pipeline already used in other studies^6; 7^ equipped with the Burrows-Wheeler aligner tool (BWA)^8^, SAMtools^8^, PICARD, the Genome Analysis Toolkit (GATK)^9^ and Annovar^10^. Briefly, sequenced reads were aligned to the GRCh38 reference human genome with an average coverage of 100x and coverage percentage of RefSeq coding region of 97.6% at 30x. Variant filtering was performed with VariantMaster^11^ for de novo, recessive, X-linked, and dominant mode of inheritance.

### Sanger validation and search for mosaicism

The primer pair to amplify the sequence containing *PAK2* c.1303 G>A, p.(Glu435Lys) was designed using program Primer3 (http://primer3.ut.ee) and checked for SNP absence with the tool SNPCheck V3. The forward oligonucleotide primer sequence is 5’TGGTTACACGGAAAGCTTATG3’ while the reverse sequence primer is: 5’TTGAACAAGGCTGGTGTTTAACT3’. An amplicon of 141 bp is amplified.

PCR conditions were set up by using a DNA control (NA12877, Coriell Institute). The polymerase GoTaq G2 Hot Start M743A of Promega was used in the recommended conditions. The PCR amplification program was: 94°C, 3min / 14x (94°C, 45sec / 63°C dT-0.5°C per cycle / 72°C, 45sec) / 35x (94°C, 45sec / 57°C, 30 sec / 72°C, 45 sec) / 72°C, 3min / 10°C, inf. The specificity of the primer pair was checked by agarose gel electrophoresis; a single band at the expected size was observed followed by a sequencing performed by Microsynth AG. PCR products were analyzed for ultra-deep high throughput sequencing in a NovaSeq (Illumina). The coverage was >10000 reads per sample.

### Structural modeling

The PAK4 kinase domain structure, stored under 5UPL code in the Protein Data Bank, was used as a template for the PAK2 modeling^12^. PAK4 shares 52% of sequence identity with PAK2 kinase domain. The 50 homology models of PAK2 were calculated using Modeller 9.18 software^13^. The best model was chosen based on its best overall DOPE score^14^. In the best PAK2 model the mutation of p.(Glu435Lys) was performed 5 times with BuildModel command of FoldX 5^15^. The average difference in structural stability between the wild type and mutants was calculated with FoldX 5. The UCSF Chimera software was used for structural visualization and figures creation^16^.

### Cell transfection

Glu435Lys mutation was introduced by PCR-based techniques of molecular cloning (extended primers,Forward: AAAAAAAAGCTTATGGCCCTAAAGTCGACATATGGTCTCTGGGTATCATGGCTATTAAGATGGTAGAAGGAGAGC; Reverse: AAAAAACGGCCGGTTTACTTGTACAGCTCGTCCATGCCGAGAGTGAT) into *PAK2* sequence of a plasmid designed for exogenous expression of eGFP-tagged PAK2 using HindIII and NotI restriction sites present in the original pEGFP-N2-based construct^17^. The plasmids coding for the wild-type and mutated PAK2 were transfected into HEK293T cells using jetPRIME transfection reagent (Polyplus Transfection) following the manufacturer’s instructions, the cells were cultivated for 24h, and harvested for western-blot analysis. An aliquot of cell preparation was used to check the transfection efficiency using BD Fortessa flow cytometer.

### Western-blotting

Transfected cells were washed once with ice-cold HBS (HEPES – buffered saline; 20 mM HEPES, 150 mM NaCl, pH 7.1) and scrapped into Pierce IP Lysis Buffer (#87787) with freshly added protease and phosphatase inhibitors. The suspension was then transferred to a centrifugation tube and incubated for 10 min at 4 °C. Cellular debris was removed by centrifugation (15000 g/4°C/15 min), the lysate was mixed 1:1 (v/v) with 2x Laemmli sample buffer and incubated for 5 min at 95°C.

An equivalent of 20 µg of total protein was resolved on 7.5% polyacrylamide gel (18×18 cm) and transferred to a nitrocellulose membrane. The membrane was blocked for 1h in 3% bovine serum albumin and incubated for 1h with the primary antibody in PBS with 0.1% Tween-20 (PBST), at the room temperature. It was then washed in PBST six times and incubated with the corresponding HRP-conjugated secondary antibody for 1h. The chemiluminiscence signal from Clarity Western ECL Substrate (BioRad, #170 5060) was detected and analyzed using G:BOX iChemi XT-4 (Syngene).

### Material

FRAX597 (#6029) was purchased from Tocris Biosciences, dissolved in sterile DMSO to make 10 mM stock solution, and used at 10 µM final concentration to inhibit PAK kinase activity. The antibodies against total PAK2 (ab76293) and phosphorylated Ser141/144 of PAK2/PAK1 (ab40795) were purchased from Abcam, β-actin antibody was from Sigma-Aldrich (A5441).

## Results

### Family description

We have re-studied samples from 4 members of the New Zealand family (NZKS2^18^, first published in ^4^) that has been designated as a Knobloch 2 syndrome family (KNO2). This non-consanguineous family consisted of normal parents and two affected offspring. The family was studied in the Geneva laboratory in 2007 and since then no additional tests have been done, and the elusive *KNO2* gene has not been identified. In the phenotypic assessment in 2007 the older affected brother (individual NZKS2-II-1 in ^18^) had retinal detachment of the left eye, inferior retinal detachment of the right eye, and considerable vitreous condensation in both eyes. There was no encephalocele, but the patient had a distinct tuft of dark hair in the superior occipital midline. Chest X-ray showed a mild prominence of the main pulmonary arteries and extensive interstitial changes in the mid and lower zones with bronchial wall thickening and reticular nodular opacities. His cognition at 8 years of age was normal. The younger brother (NZKS2-II-2 in ^18^) had a total retinal detachment on the left eye with no light perception and anterior cortical cataract. The majority of the right eye retina was flat, with a traction band running downwards from the disc to a peripheral vitreoretinal mass. There was an encephalocele at the midline in the parietal-occipital region that was surgically removed. Like his brother, he also had interstitial parenchymal pulmonary changes on chest X-rays, suggestive of dilated lymphatics. He also had severe developmental delay at age 6 years associated with an autistic spectrum disorder (ASD), with hardly any comprehensible speech. The previous molecular analysis of the family has excluded pathogenic variants in the *COL18A1* gene; moreover, linkage to chromosome 21 markers for a recessive disease was excluded^18^.

### PAK2 « de novo » variant p.(Glu435Lys)

The computational analysis of the exome sequencing from the samples of all four available members of the family (father, mother, and the two affected siblings) has identified a *de novo* variant (not present in parental DNA) in each sibling in the gene PAK2.

The *de novo* variant was a non-synonymous substitution p.(Glu435Lys) (NM_002577.4 :c.1303G>A, chr3: 196820520, hg38). The variant was categorized as pathogenic based on the following criteria of the ACMG^19^: PS2 (de novo), PS3 (functional studies), PM2 (not found in databases), PP3 (computational prediction of pathogenicity). The variant was absent from the gnomAD database (https://gnomad.broadinstitute.org/) V2.1.1. of 282,000 alleles and V3 of 142,000 alleles from genome and exome sequences and the BRAVO Topmed2 database (https://bravo.sph.umich.edu/freeze8/hg38/) of 264,000 alleles from genome sequences.

The heterozygous variant *PAK2* c.1303 G>A, p.(Glu435Lys) was confirmed by Sanger sequencing in the two affected siblings and it was absent in the parents DNA (Figure 1A).The variant was classified as damaging by SIFT, Provean, mutation taster, primateAI, polyphen2; the CADD score was 32. Amino acid residue Glu435 is well conserved (Figure 1B) with a GERP score of 4.7.

**Figure 1.**
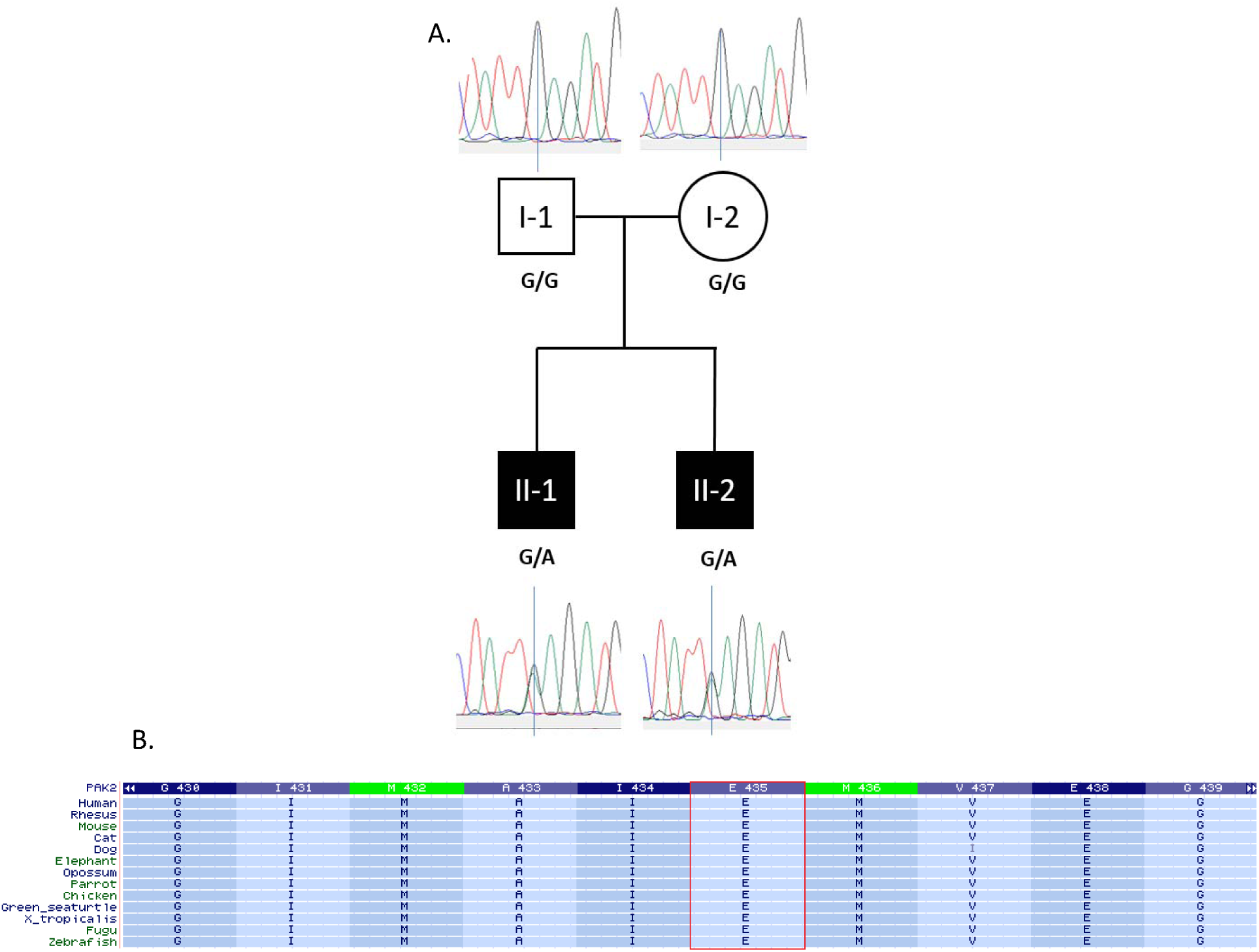
A) Family pedigree, segregation and Sanger validation of the variant PAK2: c.1303G>A. B) Glu435 is a highly conserved amino acid from human to zebrafish.

We excluded an eventual mis-paternity by calculating the percentage of high quality variants (coverage >20, GATK Quality score >200, MAF>1%) shared between I-1(father) and I-2(mother) with sons (II-1 and II-2). The sharing father-sons was 52% and the sharing mother-sons was 51.5% (Figure 2). We thus hypothesized that one of the parents has germline mosaicism potentially detectable in blood. Therefore, we PCR-amplified the genomic fragment that includes the variant and subjected the PCR product to high-throughput sequencing. More than 10000 reads were analyzed in each of the four samples (father, mother, and two affected siblings). While the variant was observed in approximately half of the reads in each affected sibling, we did not find any reads with the variant in the amplicon of the blood DNA from the parents. Therefore, we concluded that one of the parents is germline mosaic for the variant, and the mosaicism is absent from blood cells.

**Figure 2.**
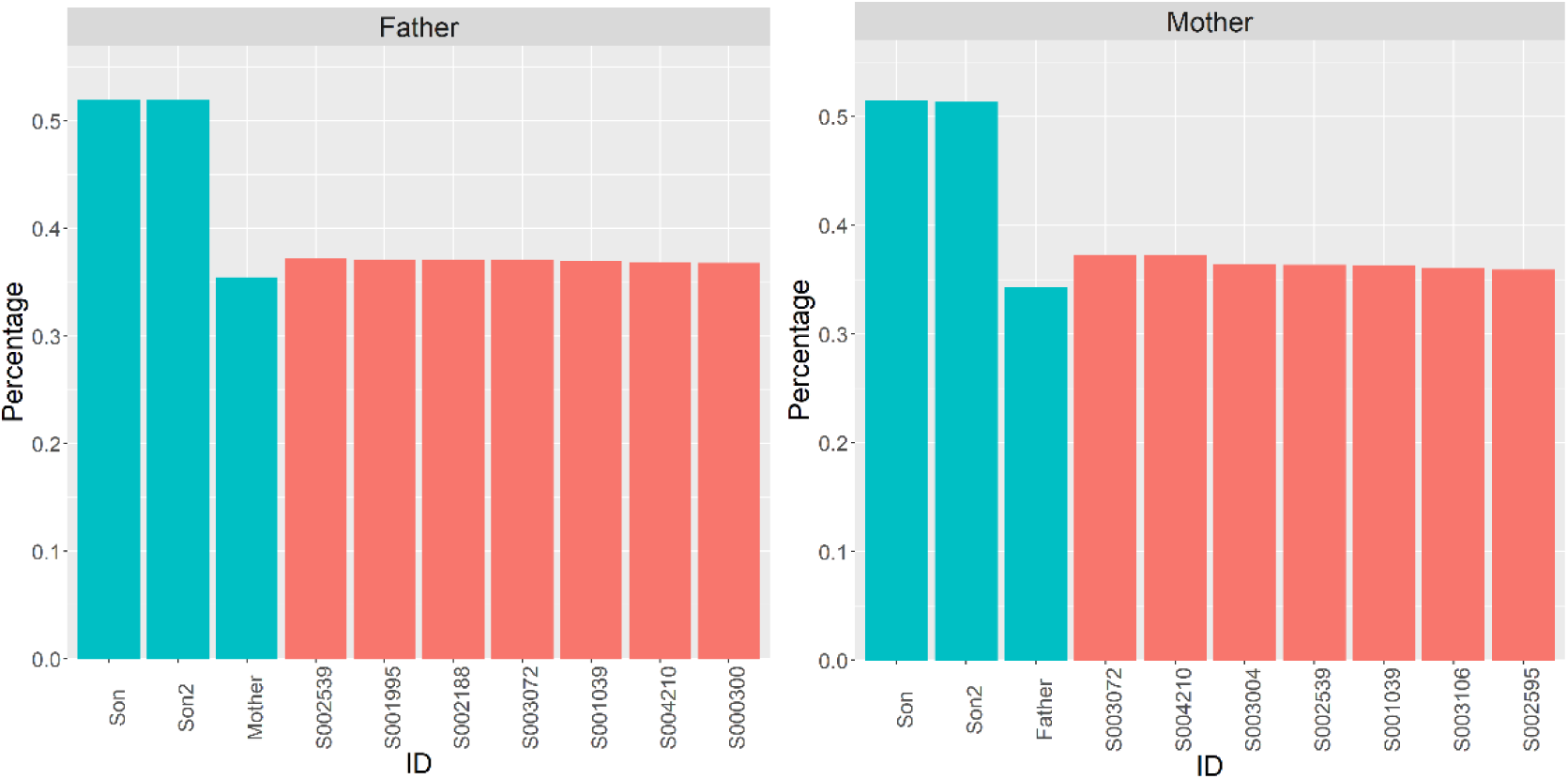
Percentage of variants (MAF>1%) shared between A) father (I-1) B) mother (I-2) and sons. In black, the sharing with 100 unrelated individuals (average father-unrelated sharing 0.33±0.026, average mother-unrelated sharing 0.32±0.027, the best 7 are represented).

### 3D structure prediction of the PAK2 p.(Glu435Lys) variant

The modeling of the PAK2 de novo variant was performed on the homology model based on the PAK4 kinase structure^12^. The p.(Glu435Lys) mutation in PAK2 is located in the center of the kinase domain, below the ATP binding pocket (Figure 3A). The Glu435 residue seems to be structurally important as it is involved in three hydrogen bond interactions: two of them with Ser371 side-chain and backbone and one to the side-chain of Trp409. The three residues are involved in the stability and conformation of helical structures (Figure 3B).

**Figure 3.**
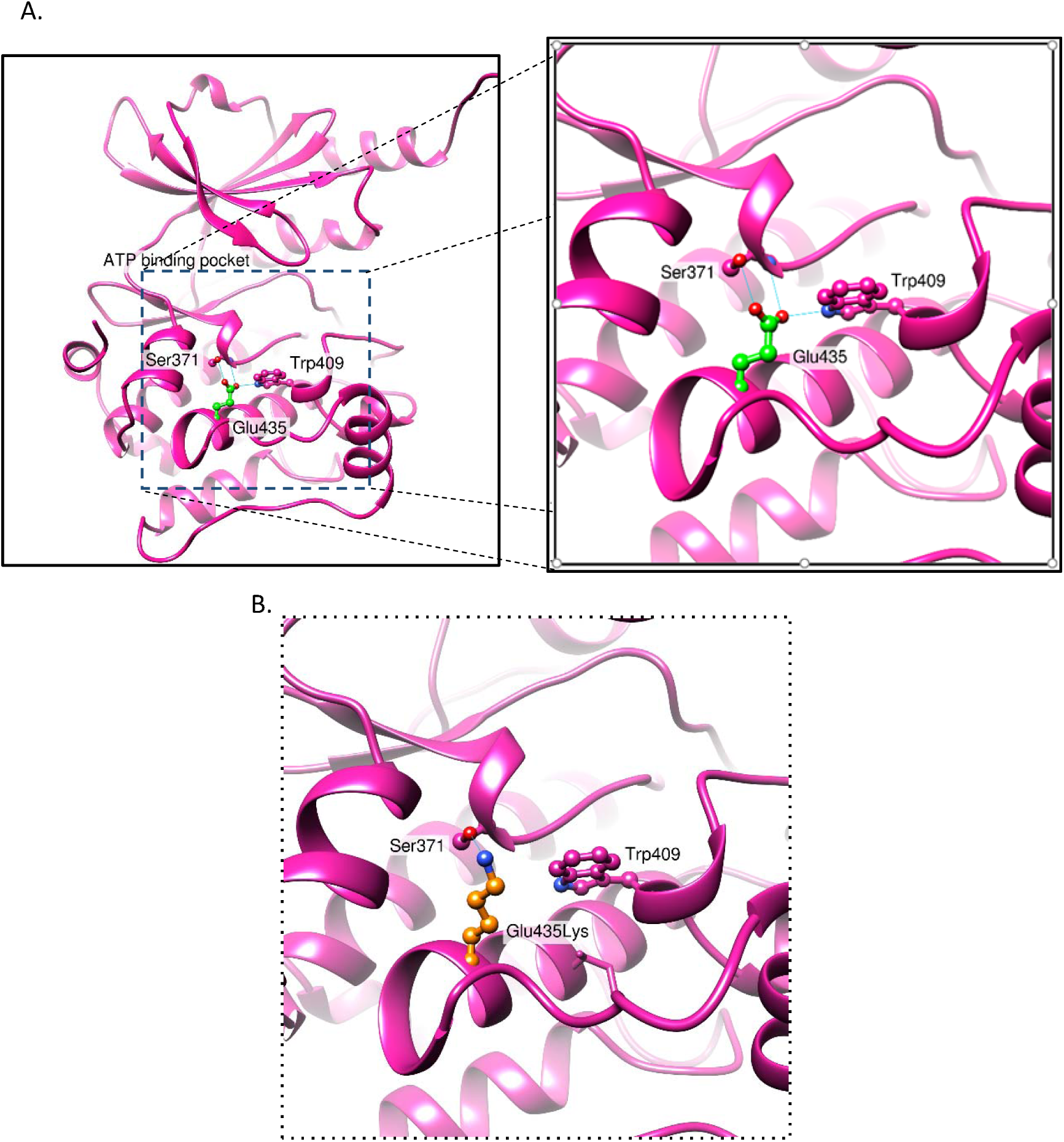
A) General structure of wild type PAK2 kinase domain model and interactions of Glu435 residue with Ser371 and Trp409 and detailed view of Glu435 residue interactions. B) Detailed view of predicted p.(Glu435Lys) mutant interactions. PAK2 is shown in ribbon representation (magenta), Glu435Lys (orange), Ser371, Thr409 residues are shown in ball and stick representation. The hydrogen bonds are shown as thin blue lines.

According to FoldX 5.0 calculations, the loss of stability upon Glu435 to Lys435 mutation is not substantial (0.26 kcal/mol with SD of 0.007), as, after the simulated mutation, p.(Glu435Lys) could still make a hydrogen bond to the side chain of Ser371. Nevertheless, the mutation most likely destabilizes the local structure, as the formation of hydrogen bond with Trp409 residue is no longer possible. The region with the mutated residue is close to the ATP binding site suggesting that the slightest changes in structure stability might affect PAK2 kinase functions (Figure 3C).

### Functional analysis of the PAK2 p.(Glu435Lys) variant

To uncover the possible impact of the p.(Glu435Lys) variant on the PAK2 kinase activity, we analyzed the extent of PAK2 phosphorylation at Ser141. This site is autophosphorylated when the kinase domain is active, and it can thus serve as a marker of PAK2 kinase activity ^12; 20^. HEK293T cells were transfected with plasmids for exogenous expression of the wild-type PAK2 (PAK2-wt) or of the mutated form (PAK2-E435K). The exogenous proteins were tagged with the green fluorescent protein (eGFP) and the transfection efficiency could thus be checked by flow cytometry: both PAK2-wt-eGFP and PAK2-E435K-eGFP were expressed in about 30% of cells, and eGFP-positive cells had closely similar mean fluorescence intensity comparing the PAK2 variants. The level of PAK2 phosphorylation at Ser141 was determined by western-blotting, using previously characterized antibodies^17^. The PAK2 p.(Glu435Lys) variant was associated with substantial reduction of phosphorylation at Ser141 of the exogenous protein, labeled as PAK2-GFP in Figure 4, panel A (pSer141/144 antibody, lane 1 versus lane 3). On the other hand, no change was induced in the total PAK2 level (panel A and B, PAK2-GFP and PAK2 endogenous). For comparison, we included samples treated with an inhibitor of PAK kinase activity, FRAX597 (Figure 4, panel A, lanes 2 and 4). As expected, the inhibitor also decreased Ser141/144 phosphorylation of the endogenous PAK2 and PAK1. We concluded that the PAK2 p.(Glu435Lys) variant results in a substantial loss of the kinase activity.

**Figure 4.**
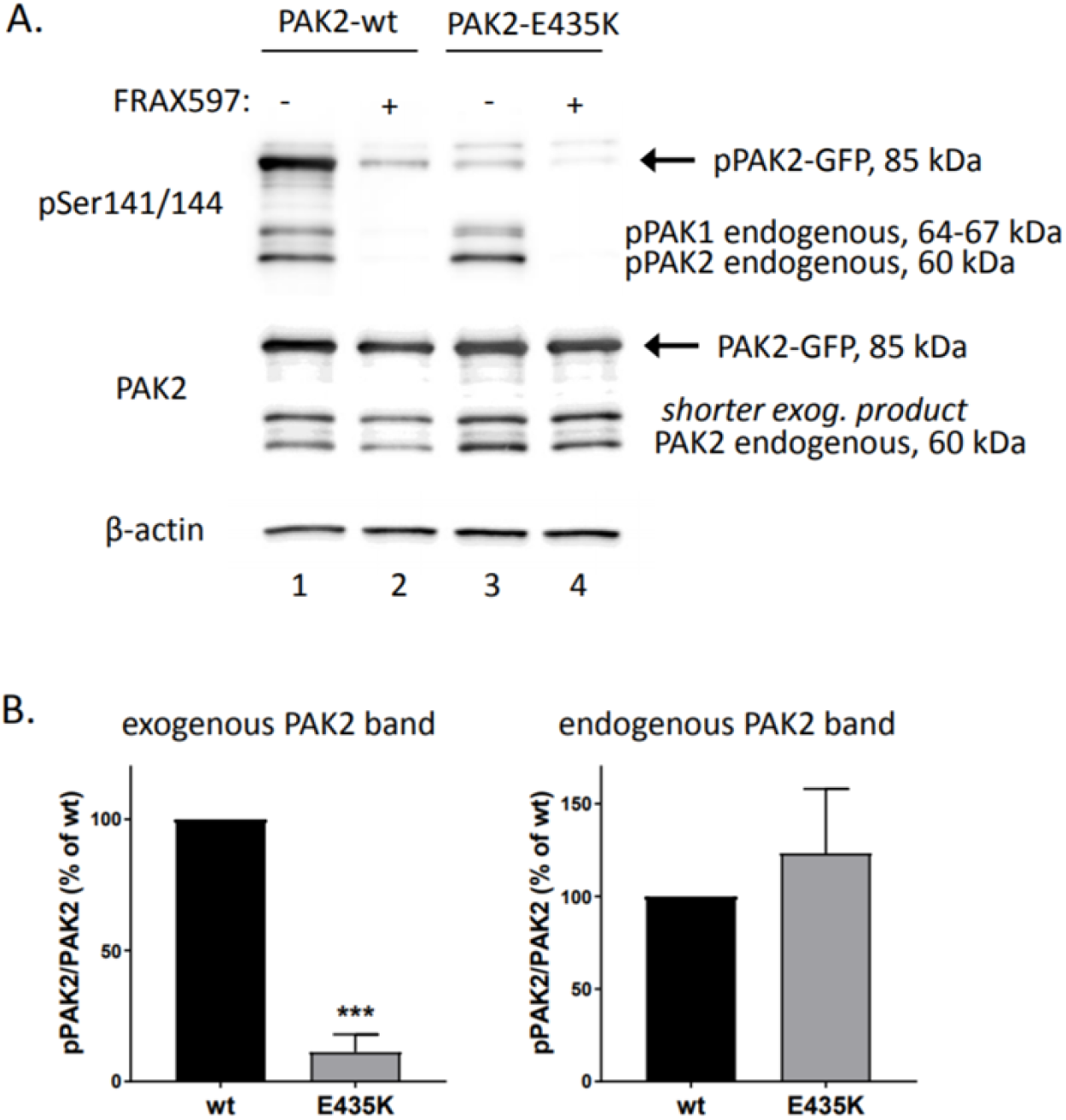
Impact of Lys435Glu mutation on Ser141 PAK2 phosphorylation. A) HEK293T cells were transfected with plasmids for exogenous expression of eGFP-tagged wild-type PAK2 (PAK2-wt) or mutated PAK2 (PAK2-E435K), and the phosphorylation status of Ser141 was analyzed using western-blotting. Some samples were treated for 1h with PAK kinase inhibitor, FRAX597, at 10 µM concentration, prior to cell lysis. B) Average pPAK2/PAK2 band intensity and standard error from four replicates. Relative band intensities from the phospho-specific antibody (pSer141/144) were normalized to PAK2-total antibody intensities.

## Discussion

### Mosaicism in de novo

The occurrence of the same pathogenic *PAK2* variants in the two affected siblings, that was absent from blood DNA of the parents, is likely to be due to parental mosaicism, i.e. a fraction of cells in the gametes of one of the parents that harbor the pathogenic variant. This phenomenon is not rare in cases of “de novo” dominant phenotypes. In a recent study of 120 apparently “de novo” cases of epileptic encephalopathy^21^, there were 10 instances of proven parental mosaicism (6 from fathers and 4 from mothers; minor allele frequency, 1.4 to 30.6%; mean number of mosaic reads after sequencing of blood or saliva DNA was 12.9%). In addition there were 2 of these 120 cases with two affected siblings in which there was an inferred mosaicism undetected in the parents. These two last cases are similar to the findings described in our study. In another study of Dravet syndrome^22^ there were 20 of 174 “de novo” SCN1A mutation cases with parental mosaicism detected in blood DNA (13 from fathers and 7 from mothers). In the years before high throughput, sequencing mosaicism has been extensively discussed. For example parental germline mosaicism has been documented in up to 5% of mothers of patients with Duchenne muscular dystrophy^23^, and in 11% of mothers of patients with hemophilia A^24^. The detection of parental mosaicism is clinically relevant since it modifies the recurrence risk.

### PAK gene family related Mendelian disorders

PAK2 is a member of a small group of serine/threonine kinases; these P21-activating kinases (PAKs) are essential in signal transduction and cellular regulation (cytoskeletal dynamics, cell motility, death and survival signaling, and cell cycle progression). They were initially discovered as binding proteins of small GTPases^25^, and are effectors of CDC42^26^ and RAC1 small GTPases (see ^5^ for review). This family of protein kinases is divided in two groups: group 1 includes PAK1, PAK2 and PAK3, while group 2 consists of PAK4, PAK5 (also called PAK7), and PAK6^5^. Mendelian disorders have been identified to date for: i/ PAK1 OMIM 602590), an autosomal dominant intellectual developmental disorder with microcephaly, seizures and speech delay (OMIM 618158)^27^; ii/ PAK3 (OMIM 300142), an X-linked intellectual disability type 30 (OMIM 300558) ^28^. PAK2 is highly intolerant to Loss of Function (LoF) mutations (pLI =0.97) and to missense (Z_miss_=3.4). According to BrainSpan (https://www.brainspan.org/), PAK2 is mainly expressed during the embryonic development in most of developing cerebral structures. PAK2 expression decreases significantly after 25 pcw (post conception weeks) and reaches a plateau at 2-3 years of age (Figure 5).

**Figure 5.**
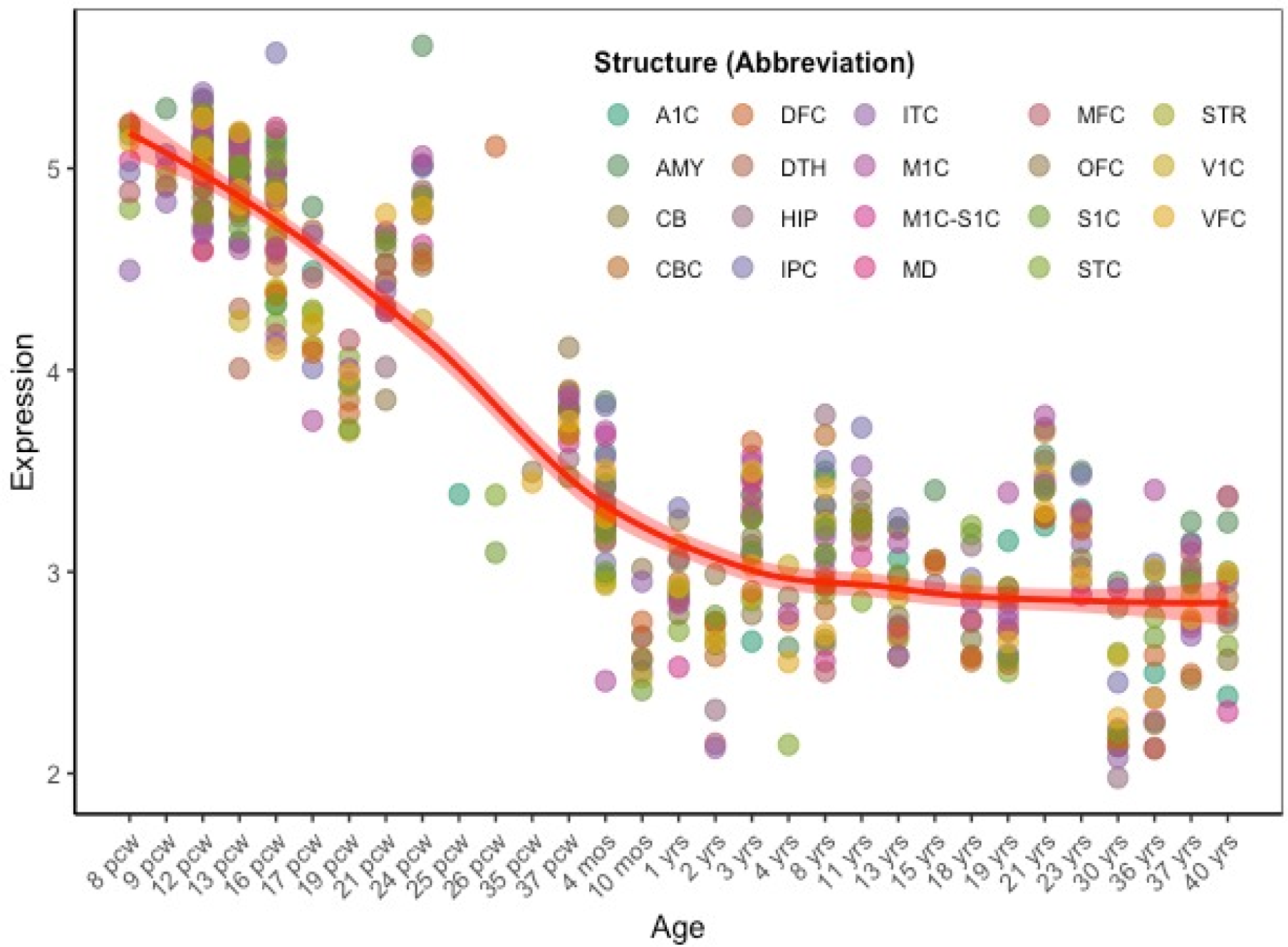
PAK2 expression (log2 RPKM) in cortical and subcortical structures across the full course of human brain development, as provided by BrainSpan. The gene is mainly expressed during the early stages of brain development; its expression decreases during pregnancy and reaches a plateau at 2-3yrs of age. For better visualization of the expressivity trend over time, the age distances have been equispaced. The red line and shaded area represent the time loess expression average across brain structures and the 95% confidence intervals respectively. pcw – post-conception weeks. Brain structure acronyms in https://www.brainspan.org

### Functional consequence of the PAK2 p.(Glu435Lys) variant

Glu435 is located within the PAK2 kinase domain. We provide evidence of a large decrease of phosphorylation at the autophosphorylation site Ser141, reflecting a loss of kinase activity^12; 20^ due to p.Glu435Lys point mutation. Thus, the acidic character of PAK2 residue 435 seems to be important for PAK2 catalytic function. Similarly, an acidic substitution of Thr402 (T402E), which is another autophosphorylation site, rendered PAK2 catalytically active^29^.

PAK group I, including PAK2, are involved in a variety of cellular processes, and mice with PAK2 knock-out do not survive an early embryonic stage of development^30^. However, some of the PAK2 functions are not kinase-dependent^31^ and might be preserved in the mutated PAK2. Presumably, considering the expression profile in early prenatal development (Figure 5), the mutation could affect cell adhesion, migration, or apoptosis, but the importance of PAK2 kinase activity for those processes is currently not fully understood.

### Knobloch syndrome

The autosomal recessive Knobloch syndrome (KNO1 OMIM 267750) has been caused by biallelic, pathogenic variants in the *COL18A1* gene^3^. Menzel et al. concluded that pathogenic variants in the *COL18A1* gene did not cause the phenotype of the family from New Zealand; thus it was proposed to name the syndrome of this family as KNO2, Knobloch syndrome 2^18^). In this study we have identified an autosomal dominant variant in the *PAK2* gene as the likely cause of KNO2. A possible functional connection might exists between COL18A1 and PAK2 since both are associated with integrin complexes (COL18A1 though proteolytic cleavage into endostatin^32^, PAK2 localizes with integrin associated adhesion complexes^17^ and is activated by RAC1 downstream of integrin matrix receptors^33^). The hypothesis that both genes are involved in the same integrin-driven pathway might explain the similarity of the phenotypic spectrum between KNO1 and KNO2.

To date, one patient with a de novo damaging mutation in *PAK2* has been reported^30^. This patient was part of a Chinese ASD (Autism spectrum disorder) cohort but, unfortunately, we were not able to obtain his complete phenotype profile. The study does not provide statistical evidence that the de novo PAK2 variant could be considered as causative to the Autism phenotype. In the same study, it was also shown that mice with heterozygous *PAK2* missense mutations display autism-related behaviors.

The family described here add to the suggestion of the implication of *PAK2* likely pathogenic variants in ASD; the phenotype of the family is mainly that of a retinal disorder and skull bone fusions defects. Therefore, at this stage the extent of the phenotypic expression of pathogenic variants in the *PAK2* gene has not yet been completed. We propose to keep the KNO2 phenotypic designation in this family for historical reasons, but to reconsider the syndrome name upon addition of more patients due to *PAK2* high impact pathogenic variants.

## Acknowledgments

The study was supported by a grant from the BLACKSWAN Foundation, Swiss foundation for research on orphan diseases (https://www.blackswanfoundation.ch), and by the Ministry of Health, Czech Republic (project for conceptual development of the research organization No 00023736 to KK and AH). We thank the Protein Modeling Facility of University of Lausanne for the bioinformatic support, and E. Ranza for assistance in the phenotypic classification. The study was partially supported by the Swiss National Science Foundation (310030_185292) and Horizon2020 (847941) to F.A.S, and the ChildCare foundation to S.E.A.

